# Characterization of the Intestinal Absorption of Morroniside from *Cornus officinalis Sieb. et Zucc* via a Caco-2 Cell Monolayer Model

**DOI:** 10.1101/2020.01.03.893768

**Authors:** Renjie Xu, Hongdan Zhu, Lingmin Hu, Beimeng Yu, Xiaohua Zhan, Yichu Yuan, Ping Zhou

**Author notes:** These authors contributed equally to this work (Renjie Xu, Hongdan Zhu). Correspondence should be addressed to Yichu Yuan; and Ping Zhou.

## Abstract

Morroniside is biologically active polyphenols found in *Cornus officinalis Sieb. et Zucc* (CO) which exhibits a broad spectrum of pharmacological activities, such as protecting nerves, preventing diabetic liver damage and renal damage. However, little data is available regarding its mechanism of intestinal absorption. Here, a human intestinal epithelial cell model Caco-2 cell *in vitro* cultured was applied to study on the absorption and transport of morroniside, the effects of time, donor concentration, pH, temperature and inhibitors, the absorption and transport of morroniside were investigated. The bidirectional permeability of morroniside from the apical (AP) to the basolateral (BL) side and in the revese direction was studied. When administrated was set at three tested concentrations (5, 25 and 100μM), the P_app_ value in AP-to-BL direction was ranged from 1.59 to 2.66×10^−6^cm/s. In the reverse direction, BL-to-AP, the value was ranged from 2.67 to 4.10 ×10^−6^cm/s. The data indicated that morroniside transport was both pH- and temperature-dependent. The morroniside’s permeability process affected by treatment with various inhibitors, such as the multidrug resistance protein inhibitors MK571 and benzbromarone, the breast cancer resistance protein inhibitor apigenin. It can be found that the mechanisms of intestinal absorption of morroniside may involve multiple transport pathways like passive diffusion as well as efflux protein-mediated active transport especially the multidrug resistance protein2 and breast cancer resistance protein. After CO was added, P_app_AB increased significantly by about 125.26%, it can therefore be assumed that some ingredients in the crude material promote morroniside’s absorption in the small intestine.

## Introduction

Traditional Chinese medicines (TCM) are natural therapeutic remedies that have been widely used for thousands of years (1). Morroniside (Fig 1), one of the most important iridoid glycosides, is the main active ingredient of *Cornus officinalis Sieb. et Zucc* (CO). It is a rich source of iridoid glycosides and has been used as a traditional Chinese medicinal herb for centuries (2). Various pharmacological studies have indicated that morroniside was effective in the treatment of Alzheimer’s disease (3) for protecting nerves (4), preventing diabetic liver damage (5) and renal damage (6), having beneficial effects on lipid metabolism and inflammation (7) and anti-anaphylactic activity (8). Since morroniside and its correlative plant extracts have exhibited these pharmacological effects, it would be hopeful that morroniside to be developed to some promising preparations of herbal medicinal products.

**Fig 1.**
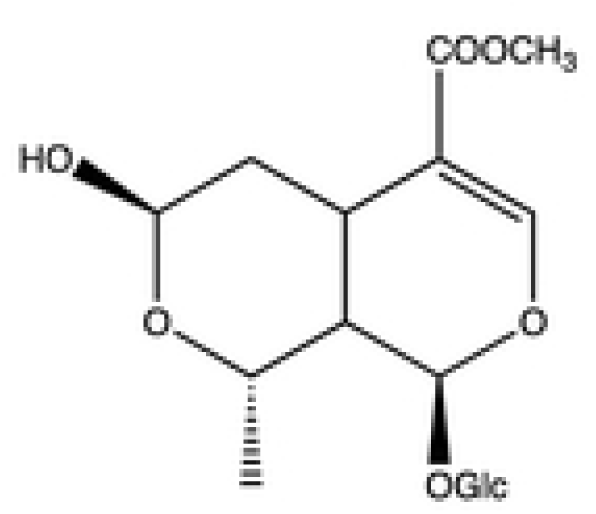
The chemical structure of morroniside.

Several studies have been conducted to determine the concentration of morroniside in biological matrices (9-11). The absolute oral bioavailability of morroniside was calculated to be 7.0, 6.1 and 3.6% for low, medium and high dosage, respectively (9, 12). It seemed like that the plasma levels of morroniside administrated intravenously were much higher than those after oral administration. It is well known that oral administration is the main route for applying TCM and they should be absorbed in the gastrointestinal tract to exhibit pharmacologic effects (13). The intestinal absorption barrier is a major factor controlling the absorption and oral bioavailability of drugs(14-16) and it is here that the first steps of pharmacokinetics occur after oral intake. Therefore, exploration of the intestinal absorption mechanism of morroniside is necessary not only for the *in vivo* pharmacokinetics study but also to provide some key information for their effective delivery system.

The aim of present study was to further investigate the intestinal absorptive characteristics of morroniside using Caco-2 cells. This model is extensively used because of its morphological and functional similarities to the human small intestinal epithelium, and it has been recognized by the FDA as a viable model that replicates human intestinal absorption (17-19). The author aimed at further revealing the reason of low bioavailability of morroniside and providing a theoretical basis for the development of formulations.

## Materials and Methods

### Materials and Reagents

Transwell permeable polycarbonate inserts (0.4 μm) and 12-well cell culture plates were obtained from Corning. (Cambridge, MA, USA). Caco-2 cell line was generously provided by Cell Bank of the Chinese Academy of Sciences (Shanghai, China). Dulbecco’s Modified Eagle’s Medium (DMEM) was obtained from Gibco Laboratories (Life Technologies Inc., Grand Island, NY, USA). Hanks’ balanced salt solution (HBSS, powder form) was obtained from Sigma Chemical Co. (Deisenhofen, Germany). Fetal bovine serum was purchased from Hyclone (Logan, UT, USA). 100× nonessential amino acids, 100× penicillin and streptomycin, 0.25% trypsin with ethylene-diaminetetraacetic acid (EDTA), 1 M 4-(2-hydroxyethyl)-1-piperazineethanesulfonic acid (HEPES), and BSA were all purchased from Invitrogen Corp. (Carlsbad,CA). Verapamil, MK 571, indomethacin, benzbromarone, apigenin, sodium vanadate and cimetidine used in this study were obtained from Aladdin Industrial Inc. (Shanghai, China). Morroniside (purity > 98.0%) and loganin (internal standard (IS), purity > 98.0%) were obtained from the National Institute for the Control of Pharmaceutical and Biological Products (Beijing, China). All other reagents (typically analytical grade or better) were used as received.

### Herb extract preparation

The extract *was* prepared according to reported methods (20). CO (100 g) crude material was decocted with 1000 mL water for 2 h. Then, the filtrate was collected and the residue was decocted with 1000 mL water for another 2 h. Finally, the two batches of filtrates were combined and concentrated to 100 mL. The solution was centrifuged to pellet any insoluble molecules then filter sterilized with a 0.22 μm (PEM) syringe filter. The contents of morroniside in preparation turned out about 16.02mg/g in crude material after determined by a HPLC method(9).

### Caco-2 Cell Culture and Cytotoxicity Assay

The conditions for Caco-2 cell culture have been described previously(21). In brief, Cells were cultured in a humidified atmosphere of 5% CO_2_ and 95% air at 37 °C. The culture medium (DMEM) was supplemented with 10% (v/v) fetal bovine serum, 1% nonessential amino acids, 1% penicillin, and streptomycin. The medium was changed every 2-3 days. When the cell monolayer reached 80-90% confluence, the cells were detached with a solution of trypsin (0.5 mg/ml) and EDTA (ethylene diamine tetraacetic acid, 0.2 mg/ml) (PAA) and reseeded at a density of 5×10^4^ cells/cm^2^.

For Cytotoxicity Assay, the cells were incubated in 96-well plate for 24 h, MTT solution was prepared at 1 mg/mL in PBS and was filtered through a 0.2 μm filter. Then, 20 μL of MTT was added into each well. Cells were incubated for 4 h at 37 °C with 5% CO_2_, 95% air and complete humidity. After 4 h, the MTT solution was removed and replaced with 150 μL of DMSO. The plate was further incubated for 5 min at room temperature, and the optical density (OD) of the wells was determined using a plate reader at a test wavelength of 490 nm.

### Transport of Analytes across the Caco-2 Monolayer

An initial stock solution of morroniside in methanol was prepared. The stock solution was then diluted with HBSS. The transepithelial electrical resistance (TEER) value was measured with a Millipore ERS voltameter (Millipore, Massa-chusetts, USA) in order to evaluate and determine the monolayer integrity. The monolayers were ready for experiments from 19 to 22 days after seeding. Only monolayers that demonstrated a TEER value above 400 Ω × cm^2^ were used for the experiment.

During incubation, the culture medium was refreshed every other day. The culture medium was discarded, 500 μL of HBSS was added to each well and incubated for 20 min at 37 °C. Afterward, each of the well was rinsed 2 times with HBSS at 37 °C. The test compounds were added to either the apical (AP, 0.5 mL) or basolateral side (BL, 1.0 mL), while the receiving chamber contained the corresponding volume of blank HBSS medium. Incubation was performed at 37°C for 180 min, with shaking at 50 rpm. To assess the drugs transport from AP to BL, at the incubation of 5, 15, 30, 45, 60, 90, 120 and 180 min, a 50 μL of the solution from BL or AP side was collected, and replaced with an equal volume of HBSS. The effect on uptake was investigated at 4 °C and 37 °C. One plate was placed in a cell incubator at 37 °C, with 5% CO_2_, and the other was in a 4 °C refrigerator. Caco-2 cells were seeded at 5 × 10^4^ cells cm^2^ in a 12-well plate. After 19 day of incubation, the effect of temperature on cellular uptake was investigated by 60 min of incubation with morroniside at 25 μg/mL. To study the effect of pH on the morroniside transport, the experiments were performed in HBSS buffer, which was adjusted to pH 6.5 or pH 7.4 on the apical side (0.5 mL) depending on the experiment, and to pH 7.4 on the basolateral side (1.5 mL).

Efflux and influx transporters were investigated for their effects on the transport flux of morroniside. Several ABC transporter inhibitors, one p-glycoprotein inhibitor (100μM verapamil) (22), three multidrug resistance-related protein inhibitor (100μM MK571, 50μM benzbromarone and 200μM indomethacin) (23-25), and one breast cancer resistance protein inhibitor (25μM apigenin) (26) were used to determine the transporters involved in the efflux transport of morroniside.; Inhibiting the influx during morroniside transport across the Caco-2 cell monolayers was investigated by adding 50μM sodium vanadate and cimetidine to evaluate the selectivity of Na/K-ATPase (27).

### Sample Processing

Samples were treated with a protein precipitation method. After spiked with 4-fold volume of acetonitrile containing 5% acetic acid and 70 ng/mL IS, the tubes were vortexed for 1min. The tubes were centrifuged at 14,000rpm for 10min, and the supernatants were collected and filtered through a 0.22μm hydrophobic membrane. An aliquot of 2 μL was injected into the LC-MS/MS system for analysis.

### UPLC-MS/MS analytical methods

An API 5500 triple-quadrupole mass spectrometer (Applied Biosystems-SCIEX, Concord, Canada) equipped with an electrospray ionization (ESI) source were used for chromatographic analyses. The separation was achieved with a ZORBAX Eclipse Plus C_18_ column (50 mm × 2.1 mm, 1.8 μm). The mobile phase consisted of 0.1% acetic acid in aqueous solution as solvent A and 100% acetonitrile solution as solvent B, and the flow rate was 0.4 mL/min. The following gradient elution was used: 0-1.0 min 1% B; 1.0-2.2 min 1→29% B; 2.2-2.3 min 29→95% B; 2.3-3.3 min 95% B; 3.3-3.4 min 95-1% B, and followed by a 1.0 min re-equilibration to the initial condition. The column was maintained at 30°C and the autosampler tray at 30°C.

Analytical procedures were evaluated with negative electrospray ionization (ESI) mode. Typical parameters for the Turbo Ionspray source set were: ion spray source temperature at 550 °C, ionspray voltage (IS) at 5000V; Gas 1 and Gas 2 (nitrogen) at 50 psi; CUR at 50; CAD at -2.0. Other parameters were shown in Table 1.

**Table 1.** Optimized MRM parameters for the determination of morroniside and IS.

| Compounds | Q1(Da) | Q3(Da) | DP(V) | CE(eV) | CXP(V) | Detected ion |
| --- | --- | --- | --- | --- | --- | --- |
| Morroniside | 451.0 | 243.0 | -100 | -30 | -15 | M<sup>-</sup> |
| Loganin | 435.2 | 227.0 | -100 | -23 | -15 | M<sup>-</sup> |

### Statistical analysis

UPLC-MS/MS data acquisition was performed using Analyst 1.5.2 and MultiQuant 2.1.1 software (Applied Biosystems). All of the data shown are the mean (standard deviation, n = 3. The data were analyzed by SPSS 20.0 Software. In the Caco-2 cell model, rate of transport is obtained from amount transported versus time curve using linear regression. The permeability of a compound is calculated using the following

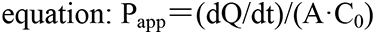

where, dQ/dt is the rate of drug transport, C_0_ is the initial concentration of the compound in the donor chamber and A represents thethe surface area of the cell monolayer. The efflux ratio P was determined by calculating the ratio of P_app_ in the secretory (BA) direction divided by that in the absorptive (AB) direction, according to the following equation:

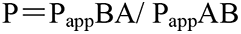

Authors had access to information that could identify individual participants during or after data collection.

## Results

### Cytotoxicity in Caco-2 Cells

Viability of cells was directly measured using MTT test to evaluate the cytotoxicity of morroniside toward Caco-2 cells prior to transport experiments. As shown in Fig 2, An increase in the concentrations of morroniside was accompanied by a slight decrease in the viability of Caco-2 cells, although the difference was not statistically significant. Even at the highest concentration (200 μM), the cell viability rates of Caco-2 cells treated with the three compounds reduced only by 8%, indicating that in our experimental design, morroniside was nontoxic to the growth of Caco-2 cells, as was 0.63mg/mL CO (containing 25μM morroniside).

**Fig 2.**
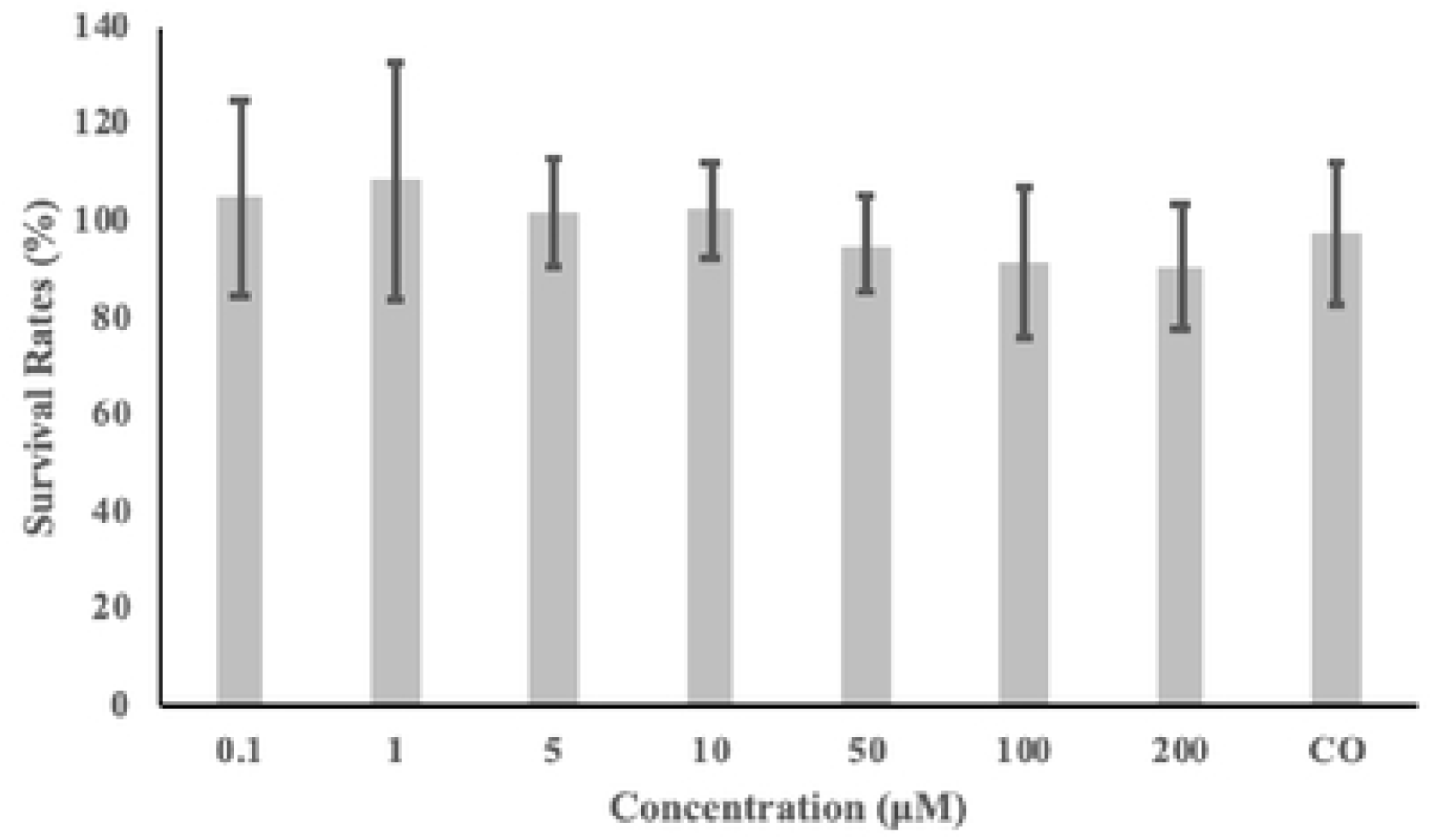
Cytotoxicity of morroniside in Caco-2 cells as evaluated by the MTT assay. Data are the mean values ± standard variation of five replications. Different letters indicate a significant difference at p <0.05.

### LC/MS/MS analysis to quantify morroniside

The retention time for morroniside and IS were approximately 1.85 and 2.01 min, respectively. As shown in Fig 3, any significant peaks interfering with morroniside or IS were not observed in fresh blank HBSS. Representative chromatograms are presented in Fig 3, including that of morroniside and IS in fresh blank HBSS and a sample 5 minutes after transport experiments. Calibration curves, constructed using linear least-squares regression, showed good linearity within a concentration range of 4-1000 ng/mL morroniside in HBSS. A typical calibration curve equation for morroniside was y=0.007x+0.0055, where y represents the ratio of the morroniside peak area to the loganin peak area (x is the concentration of morroniside). The calibration curves displayed good linearity as indicated by high correlation coefficients (R^2^=0.9997). The lower limit of quantification (LLOQ) in the collected samples was 4 ng/mL.

**Fig 3.**
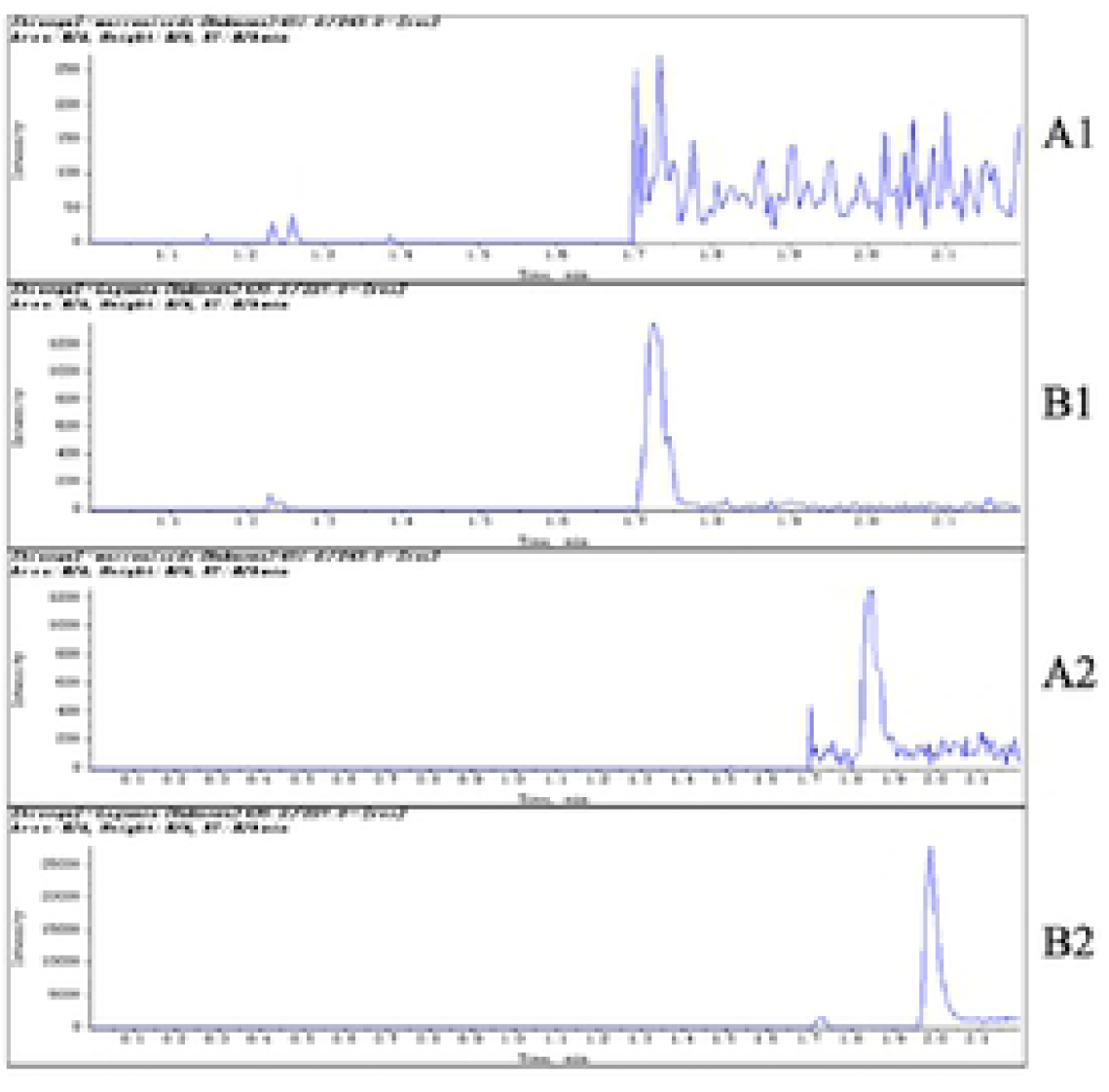
Representative multiple reaction monitoring (MRM) chromatograms of morroniside (1) and IS (2) in fresh blank HBSS (A) and a sample (B) 5 minutes after transport experiments by LC/MS/MS (7.81ng/mL for morroniside and 70ng/mL for IS)

### Uptake of morroniside by Caco-2 Cells

Results of the cellular uptakes of different concentrations (5, 25 and 100μM) of morroniside into Caco-2 cell monolayers from the AP compartment over 2 h were shown in Fig 4. As shown in Fig 4, the uptakes of morroniside into Caco-2 cells occurred quickly. P_app_AB of morroniside ranged from 1.59 at 100μM to 2.66×10^−6^cm/s at 5μM (AP to BL) and from 2.67 at 100μM to 4.10×10^−6^cm/s at 5μM (BL to AP)

**Fig 4.**
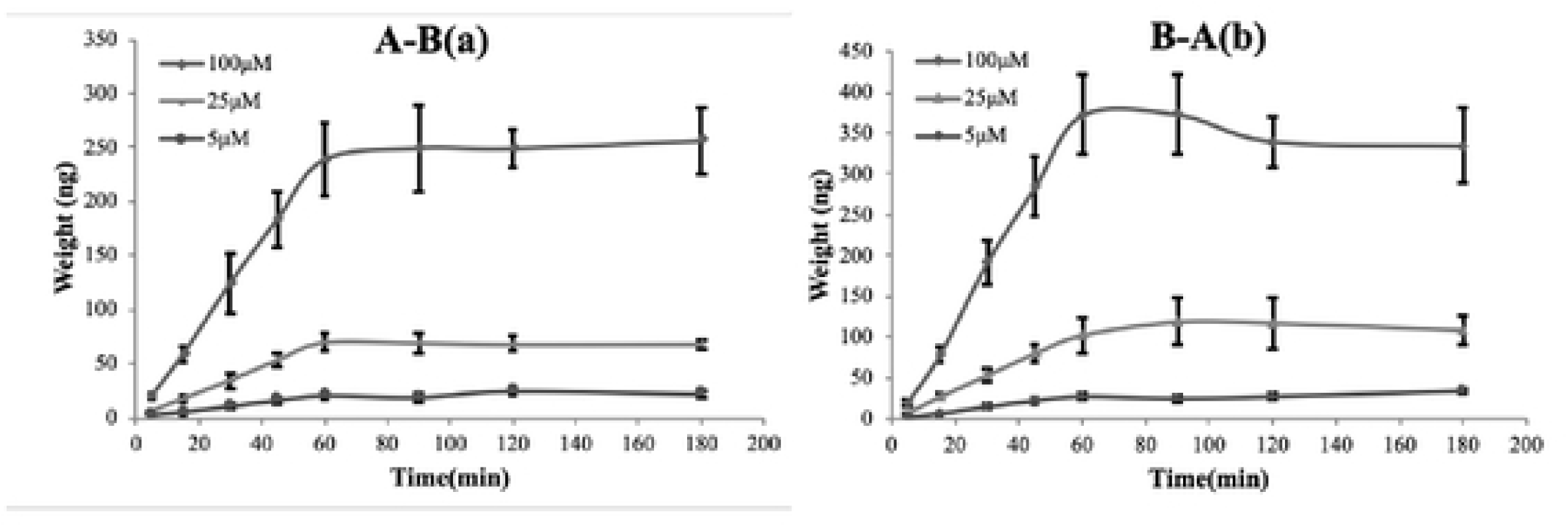
Time-course of morroniside (5, 25 and 100μM) transport across the Caco-2 cell monolayers from apical (AP) to basolateral (BL) and from BL to AP (n=3)

### Effect of pH and temperature on the permeation of morroniside

The assay studying the AP to BL transport in HBSS with various pH values was shown in Table 2. The P_app_AB values of morroniside at pH 7.4 were significantly higher than those at pH 6.0 (p<0.05), indicating an easier transport of morroniside at a higher pH (7.4) than at lower a pH (6.0).

**Table 2.** P_app_AB of morroniside (25 μM) in human intestinal Caco-2cells previously treated at different PH values and temperatures (n = 3, p < 0.05).

| Conditions | $P_{\text{appAB}}$ |
| --- | --- |
| $4^{\circ}\text{C}$ | $0.60 \pm 0.19^*$ |
| $37^{\circ}\text{C}$ | $1.94 \pm 0.64$ |
| Ph 6.0 | $1.85 \pm 0.60^{\Delta}$ |
| Ph 7.4 | $0.59 \pm 0.14$ |

\*Denote results significantly different from those of the $37^{\circ}\text{C}$ experiments
$^{\Delta}$ Denote results significantly different from those of the Ph 7.4 experiments ( $p < 0.05$ ).

Temperature plays an important role in the transport of drugs, it can not only affect the activity of the carrier protein, but also the state of drugs in solvent(28). As presented in Table 2, the morroniside transport across Caco-2 cell monolayers at low temperature (4°C) significantly reduced because P_app_AB was decreased from 1.94×10^−6^cm/s to 0.6×10^−6^cm/s.

### Effects of Various Compounds on Morroniside Transport

With the participation of 50μM of sodium vanadate or cimetidine, the uptake of morroniside did not significantly increase or decrease. These results indicated that the influx transporters like OATs or Na^+^/K^+^ pump contributed little to the transport of morroniside. (Table 3)

**Table 3.** Inhibitory effects of influx transporters on morroniside transport in Caco-2 cell monolayers.

| Inhibitors | Transporter | P<sub>app</sub>AB (×10<sup>-6</sup> cm/s) | P<sub>app</sub>BA (×10<sup>-6</sup> cm/s) | P |
| --- | --- | --- | --- | --- |
| Control |  | 1.90±0.5 | 3.06±0.6 | 1.60 |
| Sodium vanadate | Na<sup>+</sup>/K<sup>+</sup>pump | 1.89±0.4 | 3.11±0.7 | 0.78 |
| Cimetidine | OATs | 1.82±0.5 | 3.1±0.7 | 1.23 |

MRP and BCRP are present in the intestinal tract of humans, which are responsible for the transport and efflux of compounds or drugs (29). Fig 5 showed the P_app_AB values increased, but had no effect on the P_app_BA, the data indicated that morroniside was effluxed by MRP.

**Fig 5.**
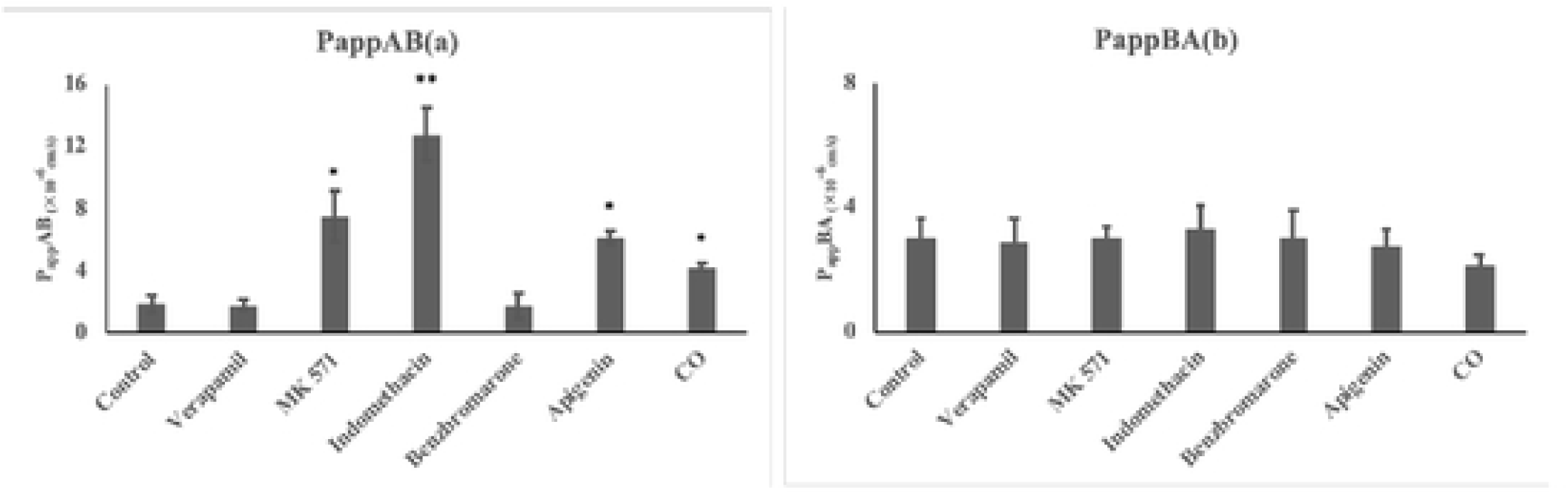
Inhibitory effects of exflux transporters on morroniside transport in human intestinal Caco-2 cell monolayers: AP to BL (a) and from BL to AP (b) (n = 3) *Denote results significantly different from those of the control experiments (p < 0.05). **Denote results significantly different from those of the control experiments (p < 0.001)

BCRP was found to be located at the apical side in the cell membrane (30). Apigenin was used to determine the effect of BCRP on the transport of 25μM morroniside. After addition of apigenin (25μM), the P_app_AB of morroniside increased significantly (p < 0.05), which resulted in a decrease in the efflux ratio by 72.05% (Fig 5). It can therefore be assumed that morroniside is also a substrate of BCRP.

## DISCUSSION

To further study the transport characters of morroniside, we investigated the relationship of time and concentration to transport. The concentration of morroniside increased approximately linearly with time from 0 to 60 min. After 60 min, the concentration gradient between the two sides had greatly decreased, result in the curves turned to reach a platform (Fig 4). A possible reason is that the concentration gradient between the two sides had greatly decreased after 60 min, which resulted in much less transport and led to a platform. From the above results, it is concluded that passive transfer is the main transport way. In our previous work, loganin (31), a structural analog of morroniside, showed good intestinal permeability by using the human intestinal Caco-2 cells model. It seemed like the cell permeability of morroniside is far lower than loganin.

Our results showed that the P_app_BA values of the morroniside was higher than their P_app_AB values (p < 0.05), indicating that some transporters might be involved in the transport of morroniside in B-A direction.

Accounting for the limit of detection of the method, compounds which are completely absorbed in the human intestine typically exhibit P_app_ values of >70×10^−6^ cm/s in the Caco-2 transwell system, whereas compounds with poor absorption (<20%) have P_app_ values of <10×10^−6^ cm/s(32). The P_app_ values of morroniside determined was at the level of 10^−6^ which indicated poor intestinal absorption about morroniside. It is also important to note that, strong metabolism may be regarded as another major cause of the low oral bioavailability of morroniside *in vivo*. However, further studies are required to clarify whether metabolism was the main factor in the *in vivo* process of morroniside.

The data illustrating how pH and temperature effect permeation of morroniside, show that morroniside transport is both pH-dependent and temperature-dependent, these results indicate, that some transporters may be involved in the efflux of morroniside. In fact, previous research indicated that both the influx (33) and efflux transport like BCRP (34, 35) were activated at lower pH levels. It can therefore be assumed that morroniside is also a substrate of BCRP. This could be indeed verified in the next subsequent experiments.

To determine whether P-gp is involved in the transport of these compounds, their transports were studied in the presence of verapamil, a known inhibitor for P-gp. But the P_app_AB and P_app_BA of morroniside did not change significantly before and after the addition of verapamil, which indicated that P-gp might not be involved in the transport of morroniside. (Fig 5)

MRP has different isoforms, from which MRP2 is located at apical side in the cell membrane and MRP3 was the main basolaterally localized MRP(36). Indomethacin and benzbromarone were used as MRP2 and MRP3 inhibitors, respectively. The results in Fig 5 showed that P_app_AB values caused a significant increase in the AP to BL flux (p < 0.001) after Indomethacin was added. This in turn indicated that morroniside is a substrate of MRP2. Co-treatment with 200μM benzbromarone did not alter morroniside’s P_app_AB or the P_app_BA values, and the efflux ratio reduced only by 6.64%.

After 0.63mg/mL CO (containing 25μM morroniside) was added, P_app_AB increased significantly by about 125.26%, and P_app_BA value was no significant difference compared with the control group. Many ingredients, especially iridoid glycosides are contained in CO. To our knowledge, the most abundant bioactive iridoid glycosides, which are widely distributed in CO, are loganin, morroniside and sweroside (37). Loganin, which was found to be a substrate of BCRP, in our previous study, may reduce efflux of morroniside and lead to a lower P_app_AB in the crude material group (31). But the compounds contained in CO are complex, and more research is needed to clarify the intestinal absorption characteristics of this herb preparation.

## Conclusions

In the present study, morroniside was shown to be poor-absorbed compound in the Caco-2 cell monolayer model and the mechanisms were mainly by passive diffusion and with efflux protein-mediated active transport. Transporters, BCRP and MRPs (especially MRP2) are vital for morroniside transport in the intestine. These new findings provided useful information for predicting its oral bioavailability, pharmacokinetics, and the clinical application as well as determination of the bioactive substance basis of morroniside.

## Acknowledgments

This work was partially supported by the Cross-fund of Biomedical Engineering of Shanghai Jiaotong University (No. XH2398) and by Xinhua hospital scientific research project (No. XH2053).

